# A simulation study investigating power estimates in Phenome-Wide Association Studies

**DOI:** 10.1101/115550

**Authors:** Anurag Verma, Yuki Bradford, Scott Dudek, Anastasia M. Lucas, Shefali S. Verma, Sarah A. Pendergrass, Marylyn D. Ritchie

**Author notes:** Corresponding Author AV YB SD AML SSV SAP *MDR.

## Abstract

**Background:** Phenome-wide association studies (PheWAS) are a high-throughput approach to evaluate comprehensive associations between genetic variants and a wide range of phenotypic measures. PheWAS has varying sample sizes for quantitative traits, and variable numbers of cases and controls for binary traits across the many phenotypes of interest, which can affect the statistical power to detect associations. The motivation of this study is to investigate the various parameters which affect the estimation of statistical power in PheWAS, including sample size, case-control ratio, minor allele frequency, and disease penetrance.

**Results:** We performed a PheWAS simulation study, where we investigated variations in statistical power based on different parameters, such as overall sample size, number of cases, case-control ratio, minor allele frequency, and disease penetrance. The simulation was performed on both binary and quantitative phenotypic measures. Our simulation on binary traits suggests that the number of cases has more impact than the case to control ratio; also, we found that a sample size of 200 cases or more maintains the statistical power to identify associations for common variants. For quantitative traits, a sample size of 1000 or more individuals performed best in the power calculations. We focused on common genetic variants (MAF>0.01) in this study; however, in future studies, we will be extending this effort to perform similar simulations on rare variants.

**Conclusions:** This study provides a series of PheWAS simulation analyses that can be used to estimate statistical power for some potential scenarios. These results can be used to provide guidelines for appropriate study design for future PheWAS analyses.

## Background

Phenome-wide association study (PheWAS) has been implemented in a variety of different studies, like within the eMERGE network[1–5], using electronic health record (EHR) information that includes international classification of disease version 9 (ICD-9) code based diagnoses, laboratory test measurements and demographic information[6–8]. Other PheWAS have used data from epidemiological studies[9,10], as well as clinical trials[8,9] such as the AIDS clinical trial group (ACTG), which consist of measurements for different clinical domains like pharmacology, metabolism, virology, and immunology[11,12]. Cohorts like these with a large number of measurements for every individual have made PheWAS a practical approach when scanning over hundreds and thousands of phenotypes in a high-throughput way. PheWAS generates genetic association hypotheses for further study and provides insights through cross-phenotype associations.

Unlike genome-wide association studies (GWAS) where one phenotype is investigated in a study population, PheWAS uses a wide range of phenotypes collected for a variety of reasons for each dataset, often with minimal curation. Thus, in PheWAS, the data collected for different measurements can vary considerably in sample size, including the numbers of cases for diagnoses, depending on the rarity of the diagnosis. This makes the estimation of statistical power for PheWAS a challenge. For example, in electronic health record (EHR) data, one of the most commonly used data types to define case-control status is through ICD-9 codes; these codes provide information on disease diagnosis, procedures, and medications in the form of three-to five-digit codes. The longitudinal ICD-9 data collected over many years varies between patients due to multiple factors, such as differences in the frequency of patient visits, differences in length of records due to different start and end dates, and incomplete patient medical history. These factors generate sparseness and missing information in the data and, hence, variability in the number of cases, the case-control ratio, and the overall sample size in case-control study designs. These factors can then affect downstream association testing. Three issues exist for measures with low sample sizes: 1) low statistical power to identify or replicate genetic associations and, 2) potentially biased estimates in analyses with low sample size, and 3) an increase in multiple hypothesis testing burden through including low powered phenotypes that may not provide insights but increase the number of statistical tests.

The goal of this study is to investigate differences in the power estimates by testing different parameters commonly used in a PheWAS analysis. We investigated these parameters using a simulation approach to characterize and determine the effect of case numbers or case-control ratio in a case-control study, as well as the sample size for quantitative trait PheWAS. With our findings from the simulation analysis, we provided recommendations to consider for future PheWAS study design.

## Methods

### Simulation Design

To investigate the power of the PheWAS approach and the sample size requirements, we simulated an additive genetic model with a range of effect sizes, case-control ratios, and minor allele frequencies. The simulation design assumes no confounding effects due to the environmental exposure and covariates. Genotypes were simulated as SNPs, which were drawn according to Hardy Weinberg Equilibrium (HWE) probabilities. Since multiple disease and phenotypes were investigated in an EHR-based PheWAS, the prevalence of the diseases could differ in the study population. In our simulated datasets, we used a constant disease prevalence, which is represented as ß_0_ in the regression model and set to 0.1[13]. We represented the effect sizes in terms of penetrance functions, which is a combination of genotypes and the risk of diseases. The penetrance function is useful in estimating the probability of disease given the genotype in a specific population. It is used to assign the case disease status to samples whose genotypes are influencing the disease risk and vice versa for controls. We used a custom script written in R[14] to generate random population-based samples with genotypes in HWE and their phenotypes using different input parameters as shown in figure 1. We simulated both binary and quantitative trait phenotypes. For quantitative trait phenotype, the same penetrance function was used to generate the normalized distribution of phenotypes where genotypes of samples at the upper and lower end of the distribution are associated with phenotype values. All the samples in the simulated datasets were drawn at random, so there is no relatedness among the individuals.

**Figure 1.**
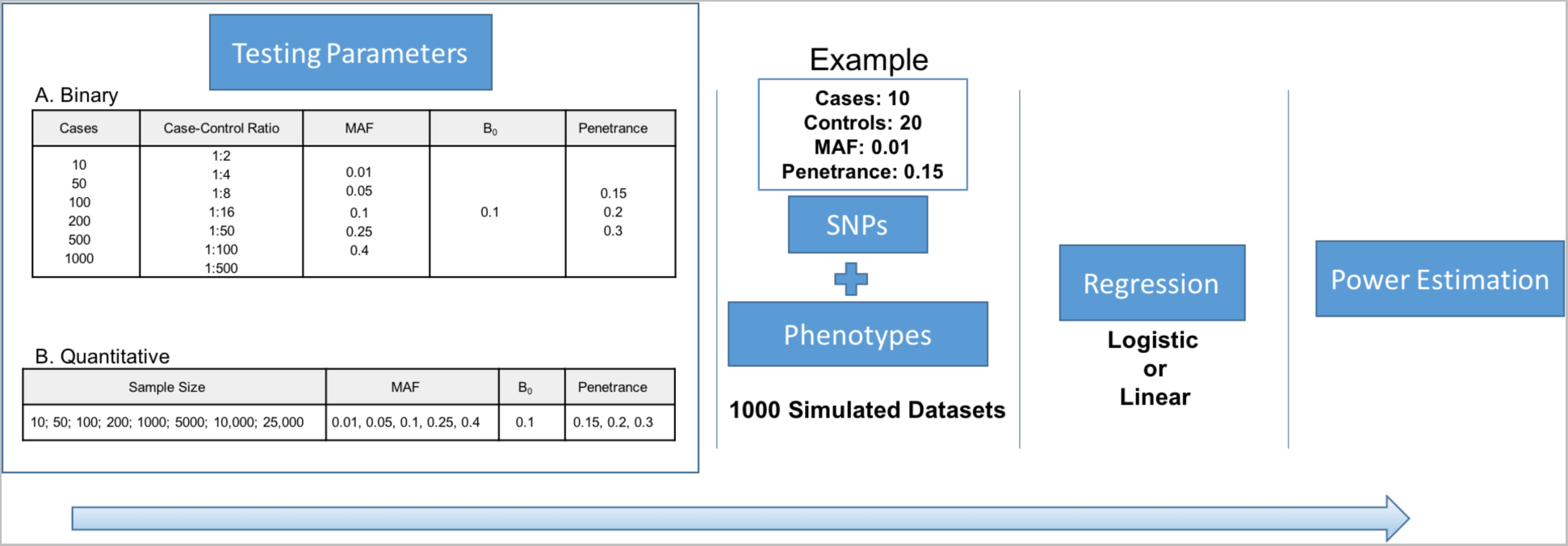
Simulation workflow. We designed a simulation approach to test different testing parameters for a PheWAS analysis and their effect on the power estimates. For each combination of the testing parameters, we generated 1,000 simulated datasets. Then, we performed association testing using logistic and linear regression for binary and quantitative phenotypes respectively. The power estimates were calculated for each combination of parameter setting at significance level of α = 0.00025 (Binary trait) and α = 0.004 (Quantitative trait)

For binary phenotypes, we generated the simulated datasets by varying the following parameter settings: cases, case-control ratios, SNP minor allele frequencies, and disease penetrance. In this simulation study, we only investigated a study population with an unbalanced and unmatched cases and controls. For example, we simulated a dataset with a random set of 30 individuals. The parameter settings for this simulated dataset were as follows: case-control ratio=1:2, cases=10, controls=20, disease penetrance =0.15, and SNP MAF=0.01. The simulated dataset was generated for four SNPs and 10 phenotypes, including one SNP-phenotype model, with signal and other models were simulated as noise. The noise was added to evaluate any systematic bias in the power estimates. The noise SNPs were generated by randomly assigning the genotypes in the study population but keeping the MAF the same as the signal SNP. We randomly assigned the cases for the noise phenotypes. Under each parameter setting, we generated 1000 datasets and then calculated associations using logistic regression. Please refer to figure 1 for all the different combinations of parameter values used for simulation.

For the continuous or quantitative trait simulations, we investigated the power estimates similarly by varying the sample size, minor allele frequencies, and disease penetrance. The simulated dataset was generated for four SNPs and one phenotype, with one signal SNP-phenotype model, and the rest was the noise data. We generated 3 noise SNPs as in the binary phenotype simulations. Again, we generated 1000 datasets for each parameter setting and then used linear regression to calculate associations with the quantitative trait. Please refer to figure 1 for all the different combinations of parameter settings used for the quantitative trait simulations. All the association testing for binary and quantitative phenotypes was performed using PLATO[15].

We calculated the power estimates by counting the number of associations below an alpha value based on total number of tests within each set of 1,000 simulated datasets for all parameter settings. For binary traits, we used α = 0.00025 (0.01/40) and for the quantitative trait, we used α = 0.004 (0.01/4).

## Results

### Binary trait simulations

We designed a simulation approach with different combinations of genotype and phenotype parameters and then performed association testing so as to investigate the factors that could influence the power to detect the signal.

In figure 2, we show trends in the estimates of power at α = 0.00025 for the parameters used in the simulations. First, we observed an increase in power with an increase in penetrance irrespective of any change in other parameters, and this is expected as highly penetrant diseases are more likely to be identified even with small numbers of samples (this is due to high effect size). We also determined that the ratio between cases and controls does not have much impact on the power. The number of cases is what primarily influences the power to detect genetic associations. For example, as shown in figure 2, the case-control ratio has a negligible effect on power, whereas with the increase in case numbers, we see the increase in the power to identify an association. These simulations also show the importance of minor allele frequency threshold when calculating associations on genotype models with an additive effect. Here, we find that all of the simulation models showed increased performance, with minor allele frequency greater than 5%. The model with lower frequency variants (MAF between 1% and 5%) did not reach 100% power until the case threshold of 1000 samples, and it was only represented in the model simulated with high disease penetrance. We observed that the common variants (MAF > 1%) have signal when there are 200 or more cases. The Type 1 error for the parameter settings used to design simulation dataset is well controlled (Supplementary figure 1) and we show an example of Type 1 error for one parameter set simulated data with cases = 200 in figure 3.

**Figure 2.**
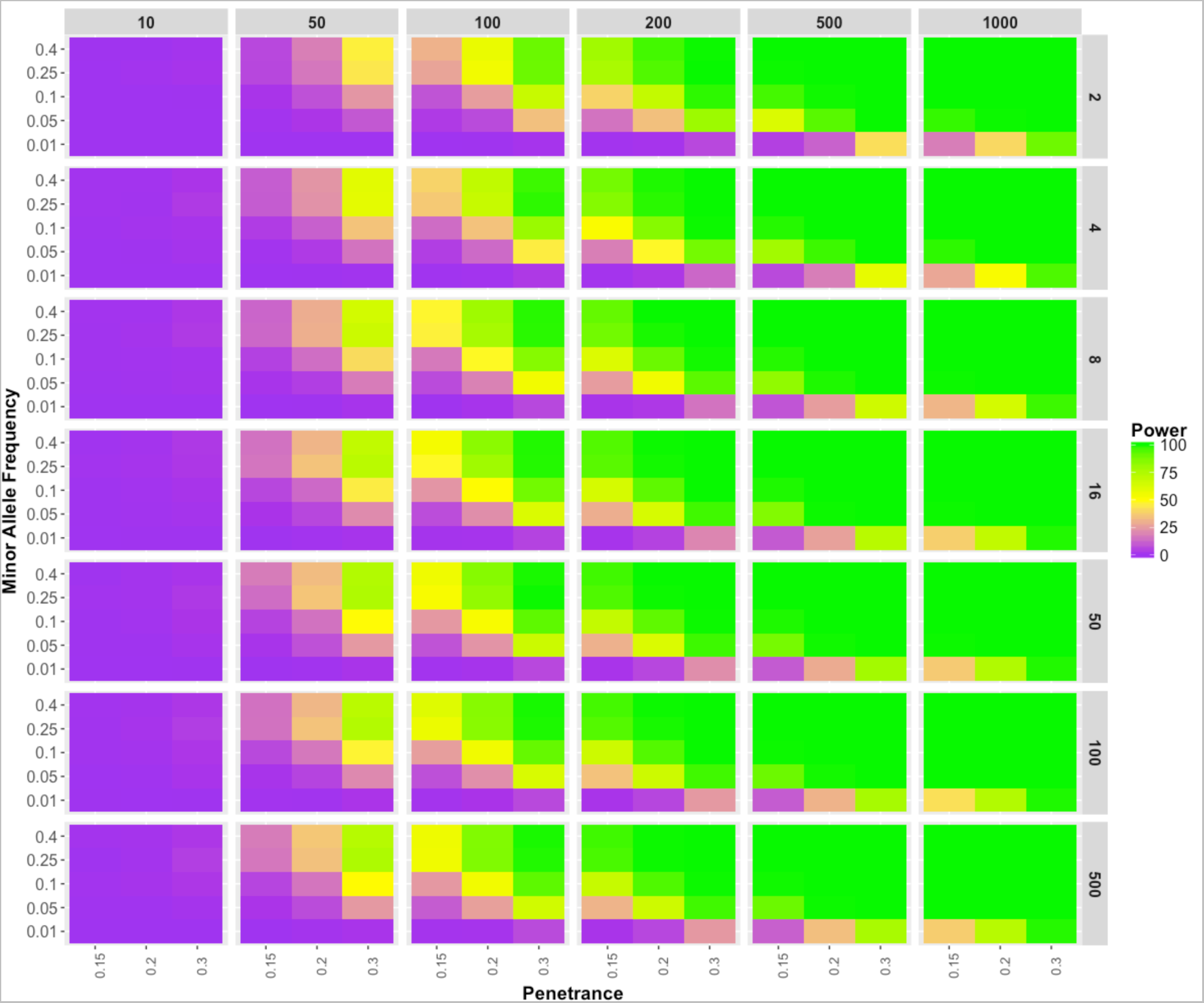
Binary Trait Power Results. The power estimates of the simulations are represented in the gradient color. Each panel represents the power estimate for a specific parameter setting. For each panel, minor allele frequencies are represented on the y-axis; the disease penetrance appear on the y-axis; the case numbers appear on the top, and the respective case-control ratios are on the right side of the box.

**Figure 3.**
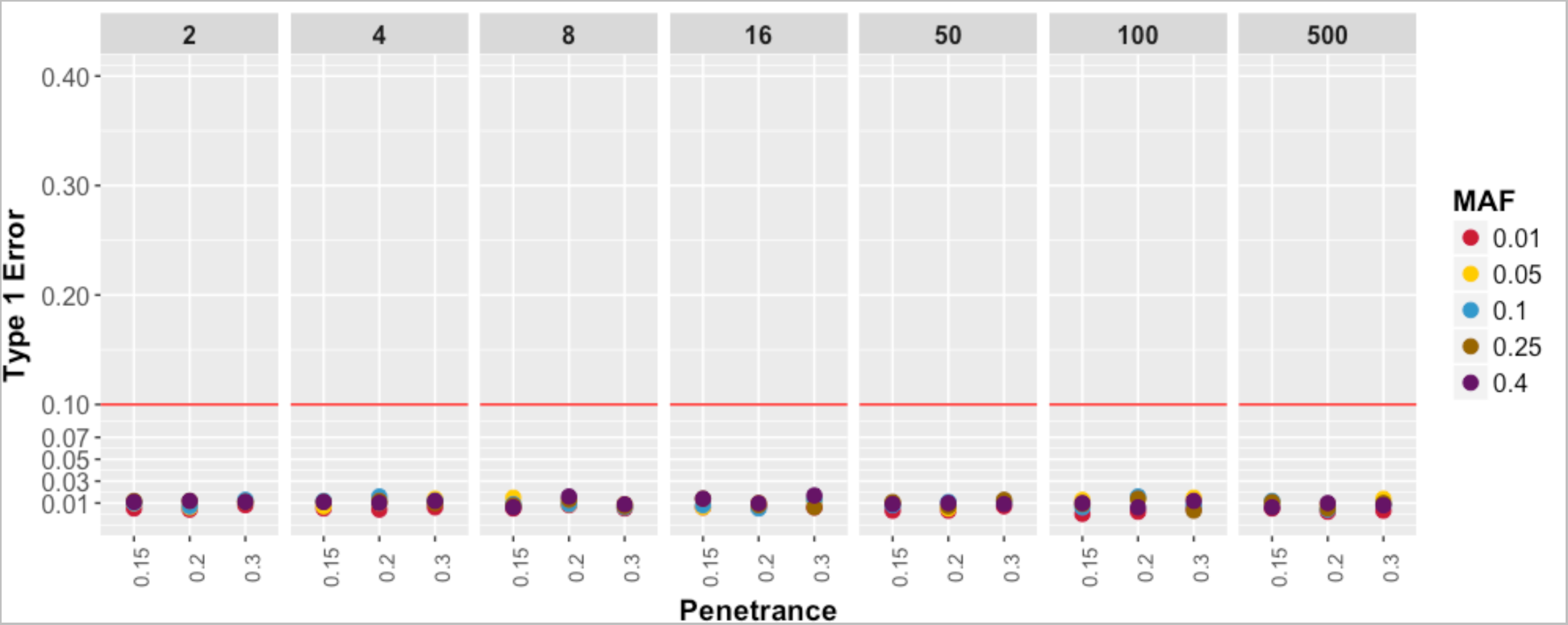
Binary Trait Type I Errors. The plot shows the Type I errors for different parameter settings at cases = 200. Each panel represents the different case-control ratio used for the simulation dataset. The Type I error on the y-axis is calculated based on the number of false positive association below significance level of α = 0.00025. The disease penetrance is represented on the x-axis and each colored point represent different MAF used in the simulations.

### Quantitative trait simulations

We also performed similar simulation analyses on quantitative traits (such as clinical lab variables) to identify a sample size threshold for multi-phenotype-based studies like PheWAS. For quantitative traits, we used different sample sizes for the simulation, ranging from 10 to 25,000, as these are based on estimates of sample sizes we observed in EHRs or clinical trials datasets[16,17]. As shown in figure 4, we observed almost no power until the dataset had approximately 1000 samples for a phenotype with a penetrance of 0.15 and; as expected, we saw the increase in power with higher penetrance, even in smaller sample sizes. Around the sample size of 1000, we see an increase in power, with a slight variation with different minor allele frequencies. Again, variants with rare minor alleles did not perform well until we had reached a sample size of 1000 and a penetrance of 0.3. These quantitative trait simulations suggest that a threshold of 1000 samples for models with MAF greater than 5% in PheWAS is recommended. Also, for the variants with MAF < 5% require either association analyses with much larger sample sizes or a different statistical approach to evaluate rare variants. As shown in figure 5, the Type I error for quantitative trait simulations are also well controlled.

**Figure 4.**
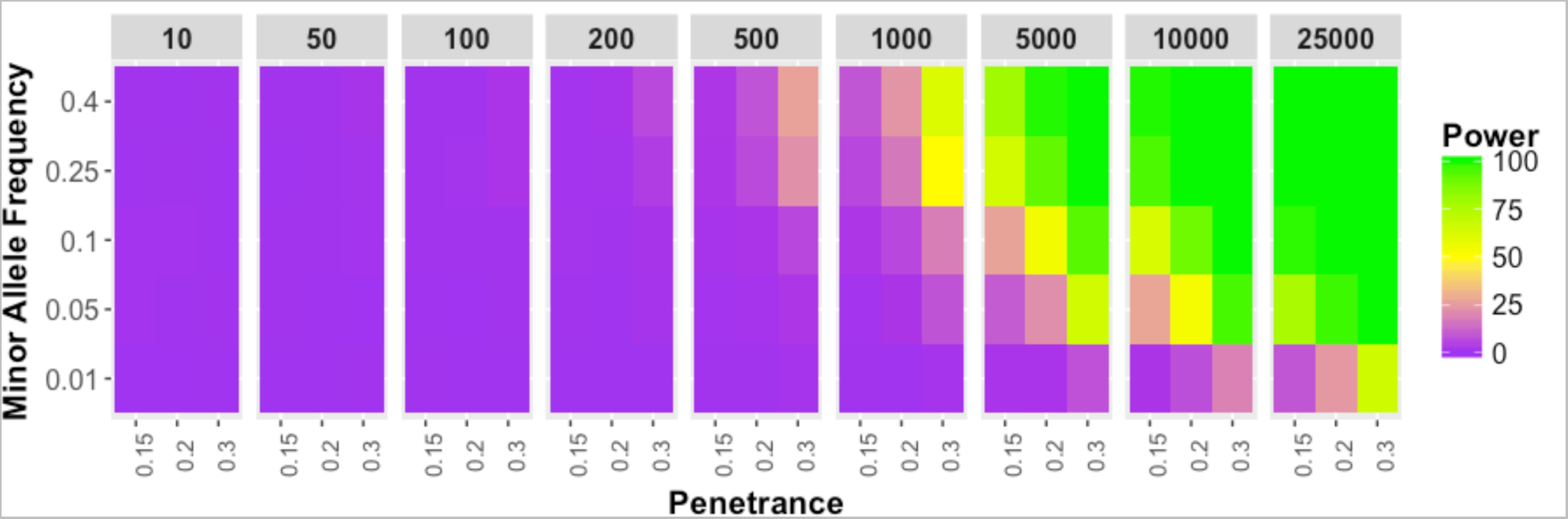
Quantitative Trait Power Results. Power estimates of the simulations are represented in the gradient color. Each panel represents the sample size of the simulated dataset, and for each panel, the minor allele frequencies are represented on the y-axis; the disease penetrance is on the y-axis.

**Figure 5.**
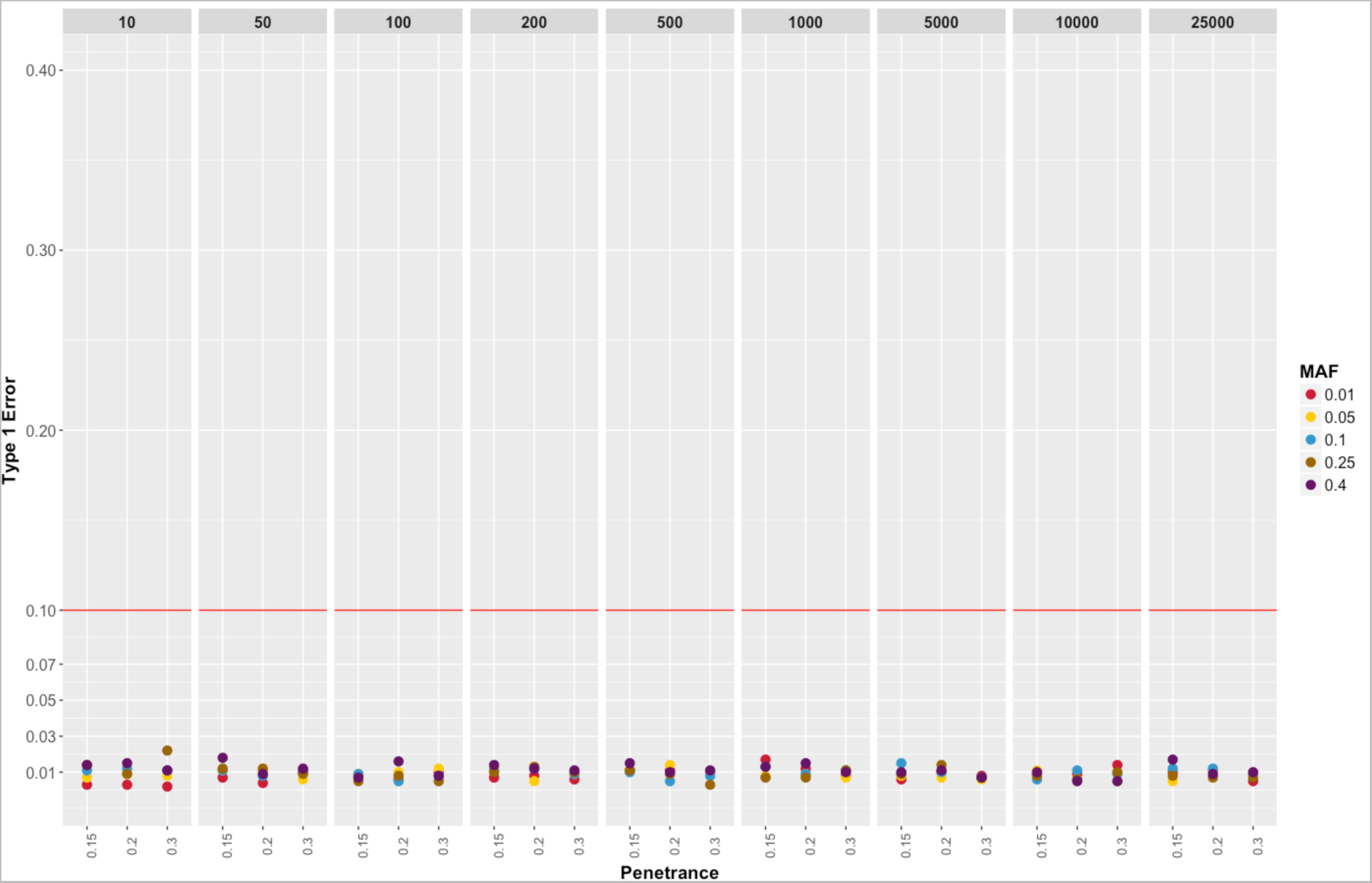
Quantitative Trait Type I errors. The plot shows the Type I errors for different parameter settings for quantitative trait simulations. Each panel represents the different sample sizes. The Type I error on the y-axis is calculated based on the number of false positive association below significance level of α = 0.00025. The disease penetrance is represented on the x-axis and each colored point represent different MAF used in the simulations.

## Discussion

PheWAS provides the genomic landscape for multiple phenotypes, but a challenge of PheWAS is the range of sample sizes and case numbers inherent when using a wide range of data instead of a single phenotype like in GWAS. For example, there are 14,025 possible ICD-9 diagnosis codes and 3,824 procedure codes used by EHR systems within healthcare provider organizations. With the introduction of ICD-10, the number of ICD-based codes has further increased to approximately 66,000. Testing 14,025 diagnosis codes for association with up to one million or more genetic variants results in a very high multiple testing burden. Usually, a large fraction of codes have very low case numbers due to the rarity of the diagnoses, and thus, they may not be sufficiently powered for association detection. For example, Geisinger Health System (GHS) is one of the largest healthcare providers in central Pennsylvania, with an EHR system including ~1.2 million unique patients. Using the EHR data in Geisinger for ~100,000 consented participants in the MyCode Community Health Initiative[16], we evaluated the extent of the variability in the number of ICD-9 codes by case count. In order to account for misdiagnosis, we defined a patient as a case for an ICD-9 code only when a patient had three or more independent visits where that specific code was represented in the patient’s record. Out of 14,025 codes for data collected between the years 1996 and 2015 for ICD-9 codes alone, 33% were not present at all and ~30% ICD-9 based diagnoses had less than 10 patients with that code (case count < 10). In figure 6, we show the trends on ICD-9 codes with cases at different thresholds and even after dropping out more than 60% of the ICD-9 codes, there are still 3,568 ICD-9 codes with 10 or more patients labeled as cases. This can increase the number of ICD-9 code based phenotypes for association testing, and it adds to the multiple hypothesis burden. In our binary trait simulations, we show that we need 200 or more cases to have enough power to detect genetic associations for common variants (MAF > 0.01). So, at that threshold, there are 831 ICD-9 codes with at least 200 patients with status as the case (as shown in figure 2). Based on our simulations, we recommend using 200 as a case threshold for the common variant PheWAS analysis.

**Figure 6.**
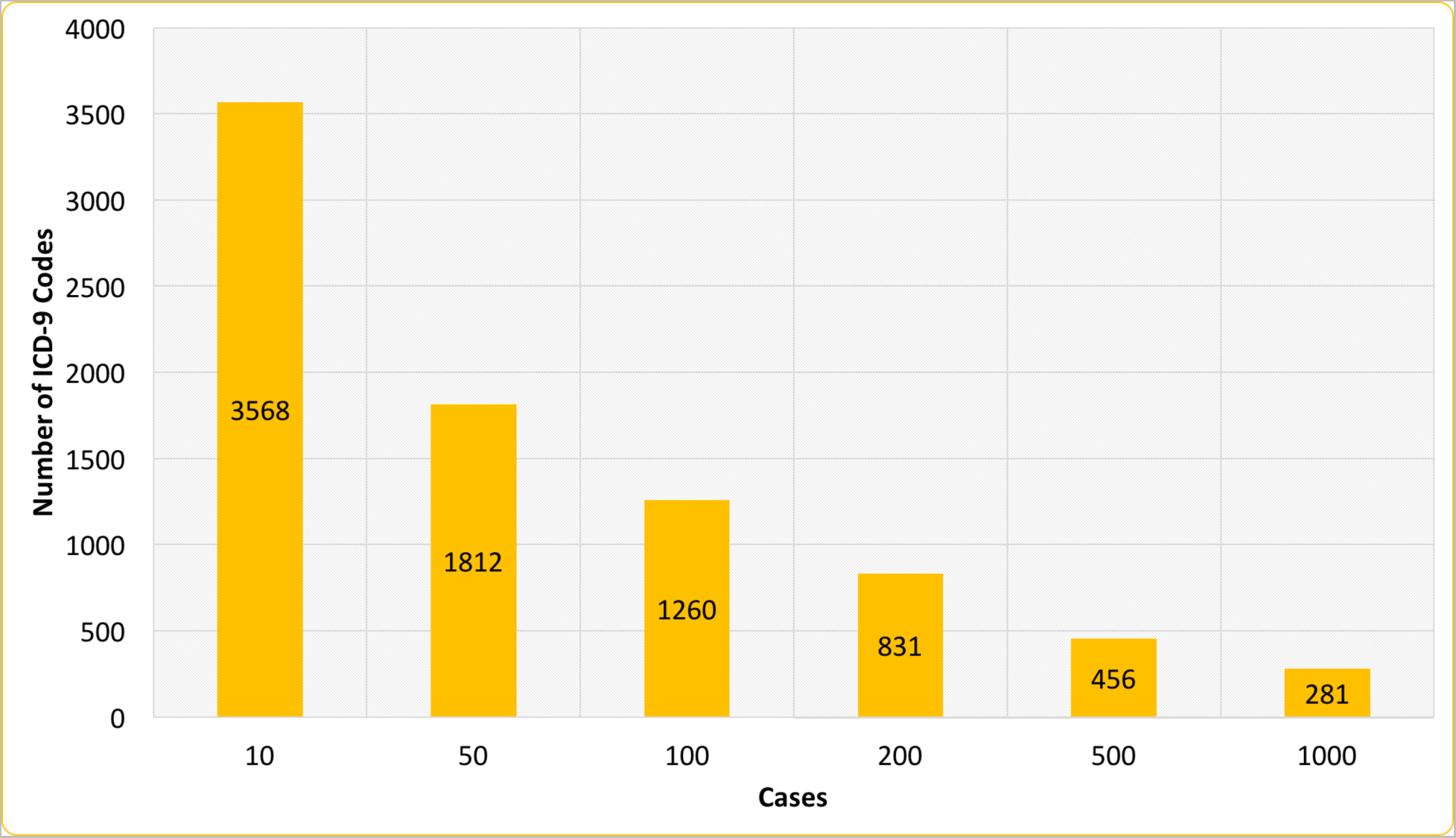
A number of ICD-9 codes, by case threshold, are generated from EHR data of 100,000 Geisinger’s MyCode participants. The x-axis represents the different case thresholds we used in the binary data simulations. The y-axis, it shows the number of ICD-9 codes for each case threshold (>=cases) in 100,000 MyCode participants. For each ICD-9 code, the cases are defined as the individuals with three or more visits to the clinic.

Using the findings from these simulations, we addressed three issues related to low sample size and its impact on PheWAS approach. First, the impact of low sample size is evident in quantitative trait simulations, which suggests that the sample size of 1000 individuals for each phenotype is important to consider in the study design. However, in binary trait simulations, we observed that overall sample size does not affect the power, but instead specifically the number of cases that drives power estimates. Secondly, low Type 1 error across all parameter settings (Figure 3, Figure 5, Supplementary Figure 1) shows no systematic bias in the regression method. However, low sample size or low case numbers will not have enough statistical power to detect the associations. Lastly, we demonstrate that using the above-suggested thresholds of case numbers for binary traits and sample size for quantitative traits can help with the selection of phenotypes and reduce the number of tests and; hence, this can reduce the multiple hypothesis testing burden.

## Limitations

Using the simulation approach, we were able to identify the parameters impacting the power to determine genetic associations and we provided recommendations for PheWAS analysis design. However, there can be other factors that can influence the power of PheWAS analysis. We primarily ran all the simulations based on a regression model (linear or logistic regression), but there are now many other statistical methods for phenome-wide association analysis[18]. Further extensions of these simulation studies to explore other statistical methods will be important. We limited our investigation in this study to an additive effect of genotypes. However, there are other factors that can influence the power estimates; these include environmental exposure, confounding covariates (age, sex, and ancestry), and underlying genotype effects (dominant and interaction). In the future, we will plan to include such effects in our simulation design.

## Conclusions

PheWAS have become a common tool to explore the genotype-phenotype landscape of large biobanks linked to comprehensive phenotype/trait data collections as in EHRs, clinical trials, or epidemiological cohort studies. This high-throughput analysis approach has been met with much success in recent years[4,6,10]. However, the community has been lacking guidance for making study design decisions regarding sample size, case to control ratios, and minor allele frequency thresholds. At present, there is not a PheWAS Power Calculator available to researchers. Thus, we implemented a large-scale simulation study to provide some guidelines for understanding the statistical power of PheWAS analyses under different scenarios. We believe these simulation results provide the needed power estimates for future PheWAS analysis decisions.

## Declarations

### List of abbreviations

PheWAS: Phenome-Wide Association Studies
MAF: Minor Allele Frequency
ACTG: AIDS Clinical Trial Group
EHR: Electronic Health Record
ICD-9: International Classification of Diseases, Ninth Revision
GHS: Geisinger Health System

## Acknowledgements

We would like to thank Dr. Tooraj Mirshahi and Dr. Janet Robishaw for helpful discussions during this study.

## Availability of data and materials

The summary results used to generate the figures are included as supplementary files.

## Authors contributions

AV and MDR conceptualized and led the project. AV contributed to designing the analysis workflow and manuscript writing. YB and AML assisted with performing the simulation analysis. SD assisted with the computer programming requirements of the project. SSV and SAP assisted with analysis design and provided important feedback on the manuscript. All the authors read and approved the final manuscript.

## Consent for publication

Not Applicable.

## Ethics approval and consent to participate

Not Applicable.

---

**Supplementary Figure 1.**
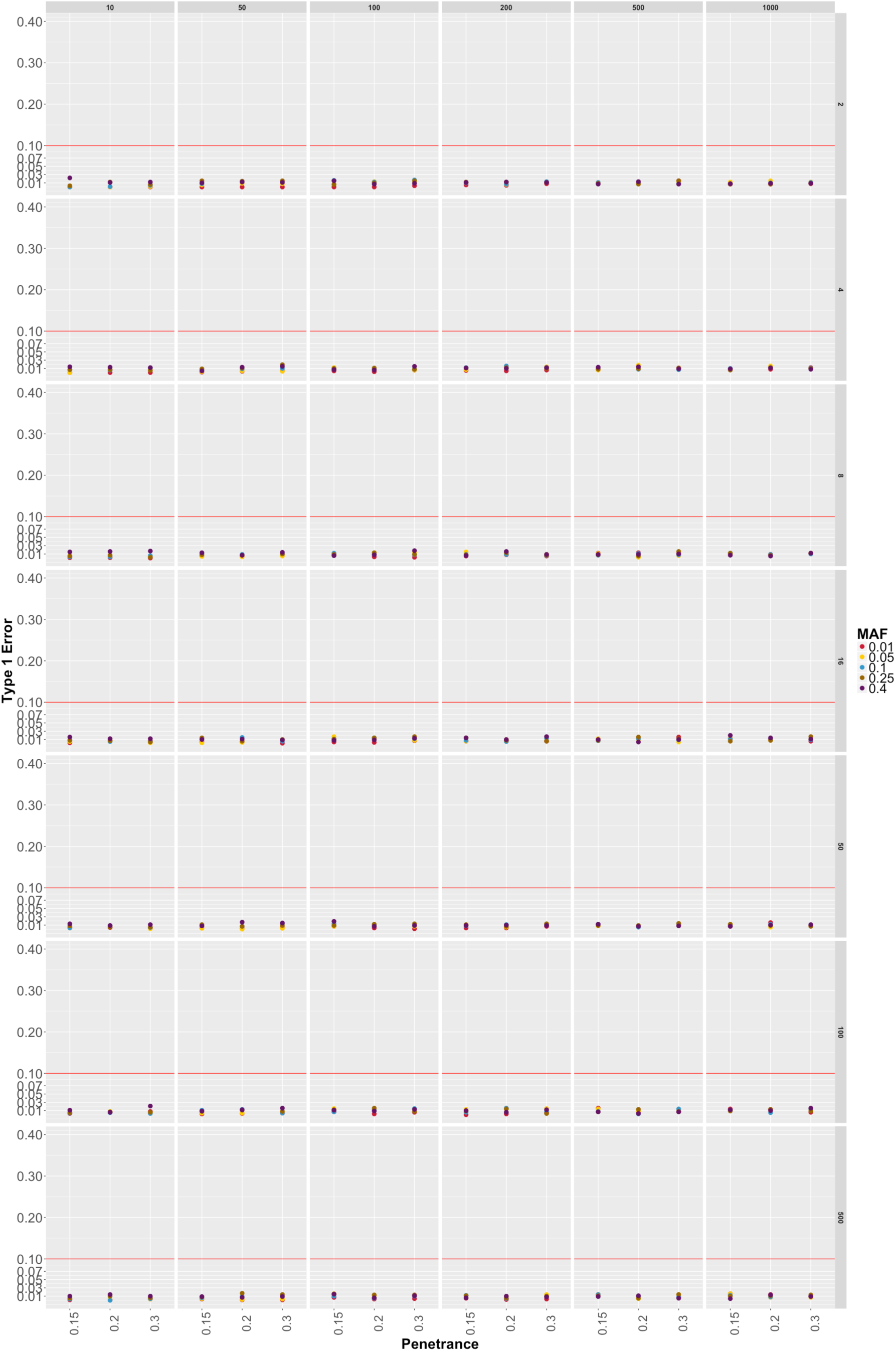
Binary Trait Type I Errors. The plot shows the Type I errors for different parameter settings. Each panel represents the different case number on the top and case-control ratio on the right which was used for the simulation dataset. The Type I error on the y-axis is calculated based on the number of false positive association below significance level of α = 0.00025. The disease penetrance is represented on the x-axis and each colored point represent different MAF used in the simulations.

